# Predicting whole-brain neural dynamics from prefrontal cortex fNIRS signal during movie-watching

**DOI:** 10.1101/2024.11.17.623979

**Authors:** Shan Gao, Ryleigh Nash, Shannon Burns, Yuan Chang Leong

## Abstract

Functional near-infrared spectroscopy (fNIRS) offers a portable, cost-effective alternative to functional magnetic resonance imaging (fMRI) for non-invasively measuring neural activity. However, fNIRS measurements are limited to cortical regions near the scalp, missing important medial and deeper brain areas. We introduce a predictive model that maps prefrontal fNIRS signals to whole-brain fMRI activity during movie-watching. By aligning neural responses to a common audiovisual stimulus, our approach leverages shared dynamics across imaging modalities to map fNIRS signals to broader neural activity patterns. We scanned participants with fNIRS and utilized a publicly available fMRI dataset of participants watching the same TV episode. The model was trained on the first half of the episode and tested on a held-out participant watching the second half to assess cross-individual and cross-stimulus generalizability. The model significantly predicted fMRI time courses in 66 out of 122 brain regions, including in areas otherwise inaccessible to fNIRS. The predicted fMRI time course also replicated intersubject functional connectivity patterns and retained semantic information about the movie content. Our publicly available model enables researchers to infer broader neural dynamics from localized fNIRS data, offering new opportunities for studying the neural basis of complex cognitive processes during naturalistic tasks.

## Introduction

Functional magnetic resonance imaging (fMRI) has become an important tool in neuroscience for non-invasively measuring moment-to-moment neural activity (Bandettini, 2020). fMRI offers the spatial resolution to study activity of specific brain regions, as well as the opportunity to measure activity across the whole brain. However, studies are constrained to tasks that can be conducted in a confined environment while participants lie still in the MRI scanner. fMRI is thus not conducive for studying many naturalistic behaviors, such as learning in a classroom environment, freely moving in a physical space, or brainstorming in groups (Varma *et al*., 2008; Shamay-Tsoory and Mendelsohn, 2019). The noise generated by the scanner and the restriction on head movement also pose additional challenges for certain populations, including children, the elderly, and individuals with sensory sensitivities (Lueken *et al*., 2011; Greene *et al*., 2016; Hausman *et al*., 2022). Furthermore, the high costs associated with fMRI scanning limits the feasibility of large-scale studies that provide sufficient statistical power and improved replicability (Button *et al*., 2013; Turner *et al*., 2018; Grady *et al*., 2021).

Functional Near-Infrared Spectroscopy (fNIRS) is emerging as a promising alternative technique for non-invasively measuring neural activity (Burns *et al*., 2019; Pinti *et al*., 2020). Similar to fMRI, fNIRS measures neural activity indirectly through the hemodynamic response (Huppert *et al*., 2006; Yücel *et al*., 2017), and prior studies have found high temporal and spatial correlation in the activation profile measured by fNIRS and fMRI during the same task (e.g., Okamoto *et al*., 2004; Cui *et al*., 2011; Sato *et al*., 2013; Noah *et al*., 2015; Liu *et al*., 2017; Wijeakumar *et al*., 2017). Relative to fMRI, fNIRS is portable, less expensive, and more tolerant of head movement. These advantages have made it suitable for a variety of studies in naturalistic settings, such as face-to-face social interactions (Suda *et al*., 2010; Hirsch *et al*., 2021), affective touch (Bennett *et al*., 2014), infant social responses (Lloyd-Fox *et al*., 2017), actors in a theater performance (Hamilton *et al*., 2018), and physical activity (Byun *et al*., 2014; Ono *et al*., 2015; Maidan *et al*., 2016). However, as near-infrared light penetrates only a few centimeters into the brain, fNIRS is limited to measuring activity in cortical regions near the scalp (Yücel *et al*., 2017). Consequently, researchers are unable to use fNIRS to measure activity in cortical regions further from the surface, such as the cingulate cortex or precuneus, or subcortical regions.

How can researchers use fNIRS to study regions beyond the cortical surface? One potential strategy is to “infer” brain activity using predictive models that map fNIRS activity onto fMRI activity in brain regions inaccessible to fNIRS by leveraging functional correlations between brain regions (Liu *et al*., 2015; Balters *et al*., 2023). Using simultaneous fNIRS and fMRI measurements, Liu and colleagues (2015) demonstrated that fMRI activity in deep brain regions could be reliably predicted from fNIRS activity measured at the scalp. More recently, Balters and colleagues (2023) “simulated” fNIRS activity by reducing the spatial resolution of cortical fMRI data to match that of fNIRS and evaluated various approaches for predicting deep brain fMRI activity from simulated fNIRS data. Their findings indicated that a linear regression model provided the most accurate predictions, suggesting a linear mapping between fNIRS and fMRI signals.

Despite the success of these predictive models, it remains unclear how well they would perform when the fNIRS and fMRI data are derived from different groups of subjects. This is a crucial consideration, as it is not always feasible to collect simultaneous fNIRS and fMRI data from the same participant. Developing procedures to train models from data collected in different groups of participants would allow us to map fNIRS data to whole-brain neural dynamics even in the absence of fMRI data from the same participant. Furthermore, previous models have primarily been trained on data collected while participants performed cognitive tasks consisting of repeated trials from a small number of experimental conditions (Liu *et al*., 2015; Balters *et al*., 2023), and may not generalize to the naturalistic tasks for which fNIRS holds the greatest promise. Indeed, naturalistic behaviors, such as having a spontaneous conversation, tap into multiple cognitive processes simultaneously and elicit a wider range of neural responses (Matusz *et al*., 2019). This simultaneous activation of diverse and overlapping brain networks may result in more complex and distributed neural patterns, making it difficult for predictive models trained on traditional tasks to generalize across different naturalistic behaviors.

The goal of this study is to develop and validate a predictive model that maps fNIRS signals to continuous whole-brain fMRI neural dynamics during movie-watching. Our study advances past work in two significant ways: first, we train a model that generalizes across individuals, addressing the challenge of applying predictive models to data from different participants; second, we train and test our predictive model on neural activity collected during a naturalistic task, movie-watching, which more closely mirrors the complexity and cognitive demands of everyday experiences. We first used fNIRS to measure participants’ neural activity as they watched a TV episode. Due to a limited number of available optodes, optodes were placed to optimize coverage of the prefrontal cortex (PFC), a decision that we had also made in our earlier work (Burns *et al*., 2019; Lyu *et al*., 2024).

We targeted the PFC due to extensive prior work implicating the region in processing complex dynamic audiovisual narratives (Baldassano *et al*., 2018; Rowland *et al*., 2018) and higher order cognition more broadly (Friedman and Robbins, 2022). Furthermore, the PFC is a heterogeneous structure composed of multiple areas that are part of distinct large-scale functional brain networks (Menon and D’Esposito, 2022). For example, the dorsomedial PFC and dorsolateral PFC are functionally coupled with the default mode network and frontoparietal network respectively. Our model can thus leverage the diverse connections of the PFC to infer neural dynamics across the brain. From a practical perspective, fNIRS signal in the PFC is often cleaner due to easier access and thinner hair coverage relative to other parts of the scalp. Thus, targeting the PFC is a strategic choice that allows us to capitalize on the region’s critical role in complex cognitive processes and its connections with major brain networks to maximize the effectiveness of accurately inferring broader neural dynamics from localized fNIRS data.

To map the PFC fNIRS data to the whole brain, we utilized a publicly available fMRI dataset where participants watched the same TV episode. We adapted a principal component regression approach to train a model that predicts whole-brain fMRI data from the fNIRS data. Importantly, the model was trained on the first half of the episode and tested on the second half of the episode in a leave-one-participant-out approach. In other words, the model was tested on data from a participant and stimuli that it was not trained on, which allowed us to assess cross-individual and cross-stimulus generalizability. To evaluate the information preserved in the fNIRS-to-fMRI mapping, we converted detailed annotations of the TV episode into textual embeddings. We then built neural encoding models (Huth *et al*., 2016; Goldstein *et al*., 2022; Caucheteux *et al*., 2023) that mapped semantic information from these embeddings onto real fMRI activity during the first half of the episode, and tested these models on fMRI activity predicted from fNIRS activity during the second half the episode. If the encoding models generalize between real and predicted activity, it would suggest that fNIRS-fMRI mapping retained information about the semantic content of the TV episode. Altogether, our study introduces a novel approach that combines the flexibility of fNIRS with the spatial coverage of fMRI, offering new possibilities for studying brain dynamics in naturalistic contexts.

## Materials and Methods

### Participants

Thirty individuals were recruited for the fNIRS study from the University of Chicago community through the research participation system managed by the Department of Psychology (SONA systems). All participants provided informed consent prior to the start of the study in accordance with the experimental procedures approved by the University of Chicago Institutional Review Board. All participants self-reported having native proficiency in English, and no hearing or speech comprehension disorders. Data from one participant was excluded because of poor data quality (see fNIRS data acquisition and preprocessing), yielding an effective sample size of 29 participants (15 females, 13 males, 1 non-binary; age: *M* = 19.69, *SD* = 0.93).

### Stimuli

Participants were scanned using fNIRS as they watched a 48min 6s segment from an episode of the BBC television series *Sherlock* (Episode 1: A Study in Pink). The stimulus was chosen due to availability of a publicly available fMRI dataset of participants watching the same segment (see fMRI dataset). The segment was divided into two runs (Run 1: 23 min; Run 2: 25 min 6s), and a short cartoon was padded to the beginning of each run to mitigate the confounding effects of stimulus-onset.

### fNIRS data acquisition and preprocessing

fNIRS data were collected using a NIRSport2 fNIRS device (NIRx Medical Technologies) with a sampling rate of 10.1725 Hz at wavelengths of 760 and 850 nm. The fNIRS device layout consisted of 20 channels composed of 8 source optodes and 7 detector optodes using the unambiguously illustrated (UI) 10/10 external position system (Jurcak *et al*., 2007). Raw fNIRS data were preprocessed in MATLAB using custom scripts that utilized the Homer2 package (Huppert *et al*., 2009). We extracted the light intensity signals from the two runs, adjusting the timecourses by 4.5 seconds to account for the hemodynamic lag. For each participant, channels were identified as noisy and removed if detector saturation occurred for more than 2 seconds or if the signal’s power spectrum exceeded a quartile coefficient of dispersion of 0.29 over the course of the scan (Dieffenbach *et al*., 2021). For each channel, the light intensity signals were converted to optical density, and filtered using a bandpass filter with a frequency range of 0.005-0.5Hz to remove physiological noises. Motion artifacts were identified and removed using targeted principal components analysis (Yücel *et al*., 2014). Motion artifacts were identified as parts of the timecourses with a signal change exceeding five standard deviations or an absolute amplitude of two within a one-second interval. A principal components analysis was then used to remove 80% of the variance in the one-second interval around the identified artifact.

Optical density signals were then converted to changes in oxygenated (HbO), deoxygenated (HbR) and total (HbT) hemoglobin concentrations following the modified Beer-Lambert law with a ppf value of 6. Data were then z-scored separately for each run and each participant. A quality assessment procedure was performed on HbO signals to identify and exclude channels that still demonstrated large signal spikes (>3SD over 1 second) after all preprocessing steps. We used the HbO time courses for all analyses because of the stronger signal amplitude, higher signal-to-noise ratio, and higher correlation to fMRI BOLD signals than HbR and HbT signals (Strangman *et al*., 2002; Tong and Frederick, 2010; Duan *et al*., 2012).

Preprocessing steps were all performed separately for each run and each participant. We then resampled each HbO time course to match the sampling frequency of the fMRI dataset (1 sample every 1.5s, see MRI acquisition and preprocessing). Participants with greater than or equal to 3 channels with missing values (i.e., 15% of channels) in either of the two runs were excluded from subsequent analysis (n = 1), resulting in an effective sample size of 29.

### MRI acquisition and preprocessing

We utilized publicly available raw structural and functional images from the *Sherlock* dataset, in which 17 participants viewed the same stimuli in two runs while undergoing fMRI. The preprocessed functional images were downloaded from the Princeton University DataSpace repository (Chen, 2016). The dataset was collected using a 3T Siemens Skyra, with a T2*-weighted EPI sequence (TR = 1,500 ms, echo time = 28 ms, voxel size = 3.0 × 3.0 × 4.0 mm, flip angle = 64°, field of view = 192 × 192 mm). Preprocessing of the functional images included slice timing correction, motion correction, linear detrending, high-pass filtering (140-s cutoff), coregistration, and affine transformation to the MNI space, and resampling to 3-mm isotropic voxels. Additional details can be found in the original publication (Chen *et al*., 2017).

### HbO-BOLD correlation analyses

For each fNIRS channel, we computed the average HbO activity across participants (group-mean HbO). To identify the corresponding location in the fMRI data, we estimated the MNI coordinates of each fNIRS channel using an anchor-based probabilistic conversion atlas (Tsuzuki et al., 2012). We then created a spherical region of interest (ROI) with a 5mm radius centered on the MNI coordinates of each channel. For each fMRI participant, we averaged the BOLD time course of every voxel within the ROI to obtain a single time course for each of the 20 ROIs. The time courses at a given ROI were then averaged across participants in the fMRI sample (group-mean BOLD).

To assess the consistency between neural activity measured by fNIRS and fMRI BOLD, we computed the Pearson correlation between the group-mean HbO and BOLD time courses at spatially matching locations. Statistical significance for each comparison was assessed using a non-parametric permutation test where the empirical correlation coefficient was compared against a null distribution generated by repeating the analysis with phase-randomized group-mean BOLD time courses, as implemented in the *nltools* package (Chang *et al*., 2024). A right-tailed *p* value was calculated for each channel location using the formula:

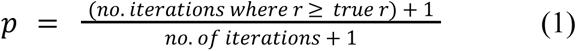

where *no. of iterations* denotes the total number of permutations and *true r* denotes the empirical correlation coefficient. Across the 20 channels, *p*-values were corrected for multiple comparisons by controlling for the false discovery rate (FDR) at *q* < 0.05 (Benjamini and Hochberg, 1995) using the *multipletests* function of the *statsmodels* package (Seabold and Perktold, 2010).

As control analyses, we extracted the group-mean BOLD time course from the primary auditory cortex, the primary visual cortex, and the average BOLD signal across the brain (i.e., global signal). The primary auditory and visual cortices were defined using spherical ROIs (5mm radius) centered around the corresponding centroids in both hemispheres in the Yeo atlas (Yeo *et al*., 2011). The group-mean auditory and visual cortices were computed by first averaging the BOLD time courses within each ROI and then across all participants in the fMRI sample. The group-mean global signal was computed by averaging the time courses across all voxels in parcels in the Yeo and Brainnetome atlases (Fan *et al*., 2016), and then averaging these time courses across participants. We computed Pearson’s correlation between the group-mean HbO time course of each fNIRS channel and (1) the group-mean primary auditory cortex BOLD time course, (2) the group-mean primary visual cortex BOLD time course, and (3) the group-mean global signal. Statistical significance was again assessed using the non-parametric permutation test described earlier, controlling for FDR at *q* < 0.05.

### fNIRS-fMRI predictive modeling

We parcellated the fMRI data into 114 cortical ROIs following the Yeo atlas (Yeo *et al*., 2011). Each cortical ROI was labeled with one of the seven cortical functional networks (visual network - VIS; somatomotor network - SM; dorsal attention network - DAN; salience/ventral attention network - VAN; limbic network - LIMB; control network - CONT; default mode network - DMN) according to Yeo *et al*. (2011). Subcortical regions (SUBC) were parcellated into 8 ROIs following the subcortical nuclei masks of the Brainnetome atlas (Fan *et al*., 2016), and included the bilateral amygdala, basal ganglia, hippocampus, and thalamus. For each participant, we averaged the BOLD time courses of all voxels within each ROI, resulting in a total of 122 time courses.

We adapted a principal component regression (henceforth aPCR; **Figure 1**) approach to predict the whole-brain fMRI BOLD time courses from their fNIRS HbO time courses. During model training, we first performed a principal component analysis (PCA; implemented with Python sklearn.decomposition.PCA) separately on the fNIRS and fMRI time courses in the training set to extract fNIRS and fMRI PCs capturing 90% of variance in the original data (average number of PCs across leave-one-participant-out iterations: fNIRS - 12.8, fMRI - 49). We then fit a linear regression model predicting fMRI PCs from fNIRS PCs.

**Figure 1.**
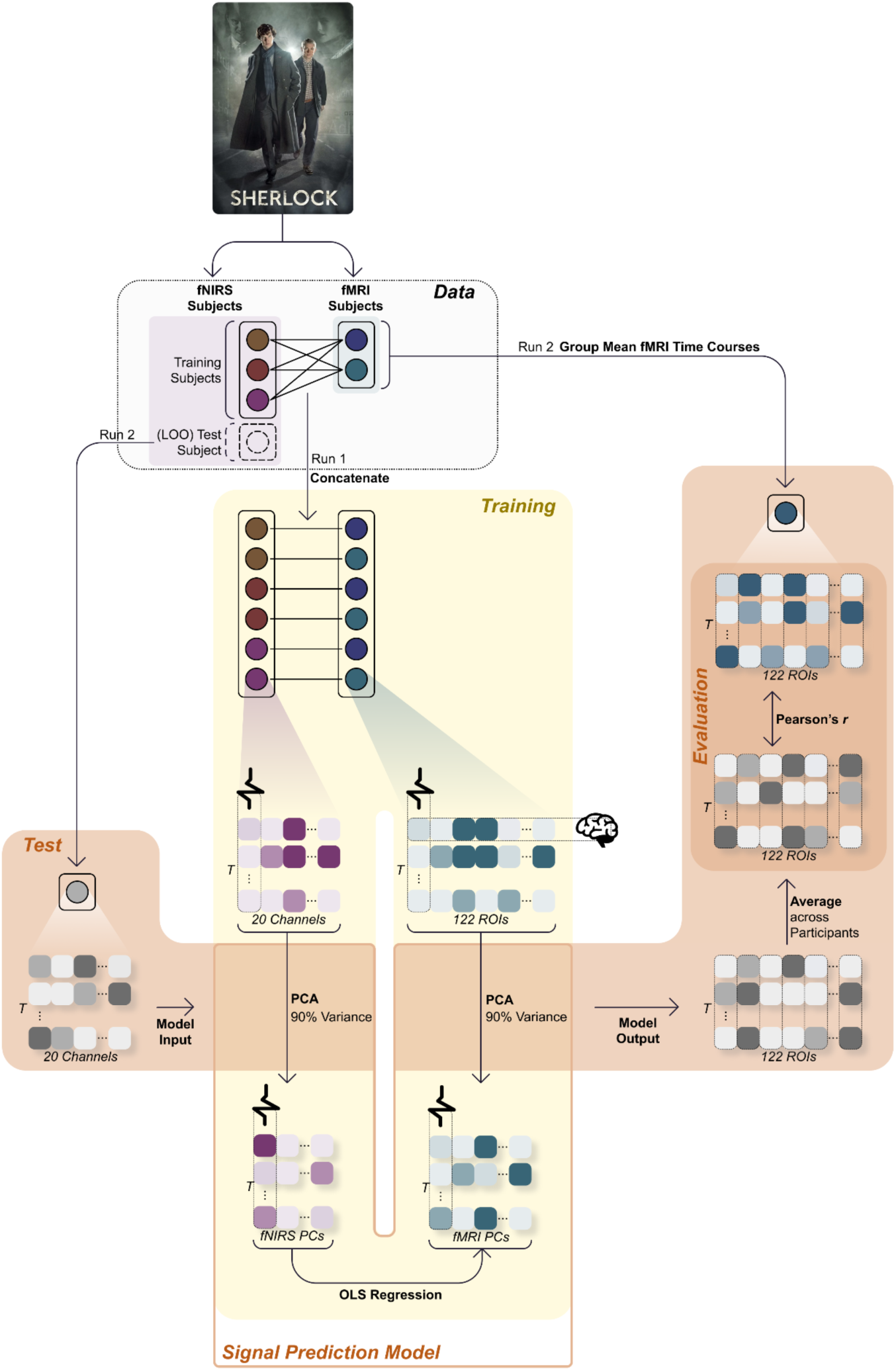
Model Training and Evaluation. The aPCR model was trained and tested following an across-run, leave-one-participant-out approach. Training involved applying separate PCAs to extract PCs capturing 90% of the variance in the fNIRS and fMRI data. Linear regression was then used to predict fMRI PCs from fNIRS PCs. Training data consisted of Run 1 data from all but one fNIRS participants and all fMRI participants. Testing data consisted of Run 2 data from the held-out fNIRS participant. Model performance was evaluated by comparing the average predicted fMRI time courses with the group mean fMRI time courses using Pearson’s correlation.

#### Train-Test split

We took a stringent across-run (train on Run 1; test on Run 2), leave-one-participant-out (LOO) approach to train and test the fNIRS-fMRI predictive model. For each LOO iteration, all data from Run 2 was excluded from the training set. In addition, all data from one fNIRS participant was held-out as the test data. Thus, the training set comprised the Run 1 fNIRS time courses from the remaining 28 fNIRS participants as predictor data, and the Run 1 fMRI time courses from all 17 fMRI participants as target data.

Because aPCR does not allow missing values in predictors or targets, we imputed missing values as follows: (1) For fMRI data, each missing value was replaced with the group mean value at the corresponding ROI and time point. (2) For fNIRS data, we first excluded channels with missing values in the left-out test participant. These channels were also excluded from the training data to match the shape of the test data. Any remaining missing values in the fNIRS training data were imputed with the group mean value across the 28 training participants at the corresponding channel and time point.

For each LOO iteration, we generated a comprehensive training set where each of the 28 remaining fNIRS participants was paired with each of the 17 fMRI participants, resulting in 28 x 17 = 476 unique pairings. Data from each pairing were then concatenated along the time axis, such that each fNIRS participant’s data was used to predict each fMRI participant’s data. This approach exposed the model to a wide variety of data pairings, improving its ability to generalize to new participants.

#### Model evaluation

In the testing phase, we applied the trained aPCR model to the Run 2 fNIRS time course of the held-out participant. In other words, the model was tested on data from a participant and watching a part of the movie that was not in the training data. For each LOO iteration, the model’s output consisted of the predicted time courses for each fMRI PC, which were then back-projected to the original ROI space using the inverse of the fMRI PCA transformation, resulting in predicted BOLD time courses for the 122 ROIs.

To assess model accuracy, we compared the average predicted fMRI time courses across the 29 LOO iterations to the average fMRI time courses from Run 2 across the 17 fMRI participants. Specifically, for each ROI, we computed the average predicted fMRI time course across all LOO iterations and then correlated this average predicted time course with the average fMRI time course from the real data. Statistical significance was assessed by comparing the true correlation values against a null distribution generated by repeating the correlation 1000 times with phase-randomized average fMRI time courses. *p*-values were computed following Equation 1 and corrected for multiple comparisons across the 122 ROIs by controlling for FDR at *q* < 0.05.

To further assess model accuracy by functional network, we grouped the 122 ROIs into the seven cortical networks defined by the Yeo atlas and one subcortical network that included all subcortical ROIs. For each network, we computed the proportion of significant ROIs and the median model performance. We computed the median correlation value rather than the mean correlation value as the former does not require Fisher transformation to normalize the distribution of correlation coefficients (Chen *et al*., 2016), and is a measure of model performance that relies on fewer assumptions.

### Inter-subject functional connectivity analyses

In addition to testing whether the model predicts ROI activity, we also examined the extent to which the model predictions recapitulated whole-brain patterns of functional correlations between ROIs. To that end, we examined inter-subject functional connectivity (ISFC), which isolates the stimulus-driven component of functional connectivity (Simony *et al*., 2016). Specifically, we compared ISFC patterns between the observed and predicted fMRI time courses for Run 2. We calculated the Pearson correlation between the time course of one ROI for that participant and the time course of another ROI across all other participants. This process was repeated pairwise for every pair of ROIs, and then repeated across all participants. We then extracted the median correlation value for each pair of ROIs. This procedure was performed separately for the observed and predicted fMRI time courses, resulting in two ISFC matrices.

The diagonal of an ISFC matrix reflects the ISC of each ROI, and was thus excluded from all ISFC analyses. Additionally, an ISFC matrix is not symmetrical along the diagonal (i.e., the correlation between region 1 and the average time course of region 2 is not necessarily the same as the correlation between region 2 and the average time course of region 1). Following Simony and colleagues (2016), we averaged the corresponding cells in the upper and lower triangles of the matrix to obtain a single value representing the ISFC between two regions. To evaluate the extent to which model predictions recapitulated ISFC patterns, we computed the Pearson correlation between the ISFC values from the predicted BOLD time courses and those from the observed BOLD time courses. Statistical significance was assessed using a non-parametric Mantel test (Mantel, 1967; Kriegeskorte *et al*., 2008), where the analysis was repeated 1,000 times while shuffling the ROI labels of the observed BOLD time courses to generate a null distribution, with *p*-values computed following equation 1.

### Semantic encoding models

We obtained detailed annotations of the movie content provided by Chen and colleagues (2017). These annotations divided the movie into 50 scenes spanning 1,000 shorter segments, with annotations describing what happened during each segment (e.g., “Sherlock picks up a small pink suitcase from a chair and brings it back into the living room”). We then processed these annotations using the Universal Sentence Encoder, converting each segment description into a 512-dimension vector embedding that captures its semantic meaning (Cer *et al*., 2018). To align these segment embeddings with the fMRI data, we resampled the embeddings to match the TR of the fMRI scans. Each TR was assigned the embedding of the segment it corresponded to. If a TR spanned two segments, the TR’s embedding was calculated as the average of the embeddings of the two segments. The segment embedding of TR *t* is denoted as SEG_t_.

The meaning of a sentence often depends on the preceding context. To capture this, we constructed a context embedding that models the accumulation of semantic information over time. This context embedding, CTX, was initialized at zero at the beginning of each scene, reflecting the reset of context at event boundaries (Zacks *et al*., 2007; Pu *et al*., 2022). At each TR *t*, CTX_t_ was updated to be the average of CTX_t-1_ and SEG_t-1_. CTX_t_ was then concatenated with SEG_t_, resulting in a 1024-dimension vector that reflects the semantic content of both the previous context and the current input. Our goal was to fit encoding models that map semantic information to observed and predicted fMRI data (Naselaris *et al*., 2011; Deniz *et al*., 2019; Caucheteux *et al*., 2023). To prevent overfitting, we followed the approach by Tikochinski and colleagues (Tikochinski *et al*., 2023; Tikochinski *et al*., 2024) to reduce the 1024-dimension vector to 32 dimensions using PCA. We refer to the resulting 32-dimension vector as the TR’s semantic embedding, or SEM.

Consistent with our leave-one-run-out testing approach, PCA was fit only to semantic embeddings from Run 1. We limited our analysis to the 66 out of 122 ROIs where the fNIRS-fMRI model had above-chance prediction performance. First, we tested whether the encoding model would generalize to the observed Run 2 fMRI time courses from the 17 fMRI participants. This provided a measure of the performance of our encoding model in capturing semantic information in the fMRI signal. Following previous work by Caucheteux and colleagues (2023), we fit a ridge regression model (alpha = 100) to predict the observed average ROI’s time course from the 32-dimension SEM vectors from Run 1. We then tested the model on the observed and predicted BOLD time course from Run 2. Accurate prediction in both contexts would indicate that the predicted BOLD responses retained semantic information about the movie content. Statistical significance was assessed by re-training the encoding model on phase-randomized observed Run 1 BOLD time courses and applying the re-trained model to the test data. This procedure was repeated a 1000 times to generate a null distribution, with p-values computed following equation 1 and controlling for FDR at *q* < 0.05.

### Visualization

Cortical brain plots were generated with BrainNetViewer (Xia *et al*., 2013) while subcortical brain plots were generated with *R* packages *ggseg* and *ggseg3d* (Mowinckel and Vidal-Piñeiro, 2020).

## Results

### Correlation between fNIRS and fMRI time courses during movie-watching

We used fNIRS to scan 29 participants as they watched a 48-minute segment of a British mystery crime drama series (BBC’s *Sherlock*). We also utilized a publicly available fMRI dataset where participants viewed the same segment while undergoing fMRI. In both datasets, the 48-minute segment was divided into two runs (Run 1: 23 min, Run 2: 25 min), with a self-timed break in between. To examine the extent to which fNIRS activity matched fMRI activity when participants watched the same movie, we correlated the group-mean HbO time courses of the 20 fNIRS channels with group-mean fMRI BOLD time courses at corresponding locations. Of the 20 ROIs, 18 showed significant correlation between the group-mean BOLD time course and the matching group-mean HbO time course (median *r* = 0.204, FDR *q* < 0.05; **Figure 2**; **Table S1**). Statistical significance was assessed using a non-parametric phase-randomization permutation test (see Methods). These results indicate that the hemodynamic responses captured by fNIRS mirrored the neural dynamics measured using fMRI as participants watched the same movie.

**Figure 2.**
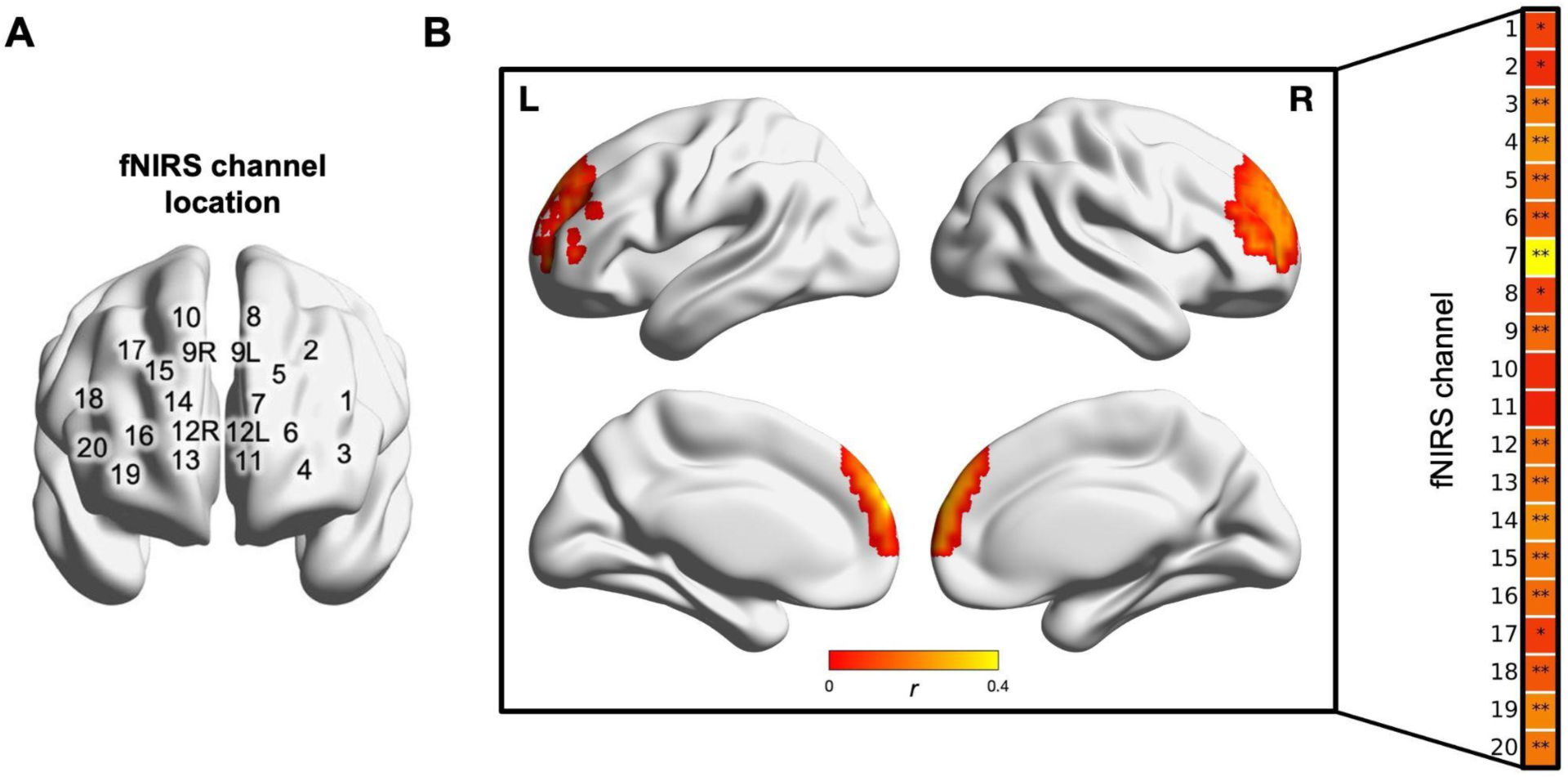
fNIRS and fMRI time courses at matching locations were correlated during movie-watching. **A.** Location of 20 fNIRS channels. **B.** Brain maps show the Pearson *r* between the 20 fNIRS channels and corresponding fMRI ROIs in the prefrontal cortex, thresholded at FDR *q* < 0.05. Statistical significance was computed using a phase-randomized permutation test. 18 out of the 20 fNIRS channels exhibited significant correlation with the corresponding fMRI ROI. **: *q* < 0.01; *q < 0.05.

As control analyses, we also computed the correlation between each group-mean HbO time course with group-mean BOLD time course at the primary auditory cortex, primary visual cortex, and the global gray matter signal derived by averaging BOLD activity of all voxels across all ROIs. We would not expect the fNIRS HbO signal in the PFC to be correlated with the fMRI signal in primary sensory regions or the global gray matter signal. Consistent with this prediction, none of the 20 channels were significantly correlated with the three control time courses (FDR *q* > 0.05; median *r*: primary auditory cortex = 0.012, primary visual cortex = 0.024, global signal = 0.028; see **Table S2** for correlation values of each ROI), indicating spatial specificity in the correlation between fNIRS and fMRI time courses.

### Predicting whole-brain fMRI signal from prefrontal fNIRS

Given the extensive and diverse connections of the PFC, we tested the extent to which we could leverage the information captured by the 20 prefrontal fNIRS channels to predict whole-brain fMRI time courses during movie-watching. Specifically, we adapted a principal components regression model to predict the fMRI BOLD activity of an independent group of participants watching the same movie (aPCR model; see *Methods*). Importantly, model training and evaluation was performed using a leave-one-out cross validation approach where the model was trained on data from Run 1 and tested on data from Run 2 of a held-out fNIRS participant. This means the model was tested on data from a participant not included in the training set, while watching a stimulus it had not previously been exposed to. This ensures that model performance did not reflect overfitting to a set of participants or a specific stimulus, and that model was generalizable to new participants and unseen stimuli.

The aPCR model significantly predicted Run 2 fMRI time courses in 66 out of the 122 ROIs (q < 0.05; **Figure 3A**). These regions included prefrontal areas covered by the fNIRS channels, including the dorsolateral PFC (left: *r* = 0.36; right: *r* = 0.29), dorsomedial PFC (left: *r* = 0.45; right: *r* = 0.44), ventrolateral PFC (left: *r* = 0.36; right: *r* = 0.36), and ventromedial PFC (left: *r* = 0.21; right: *r* = 0.27). Additionally, model performance was significantly above chance in temporal and parietal regions not covered by the fNIRS channels, including the precuneus/posterior cingulate (left: *r* = 0.29; right: *r* = 0.26), temporal parietal junction (left: *r* = 0.42; right: *r* = 0.36), intraparietal lobules (left: *r* = 0.39; right: *r* = 0.36), intraparietal sulcus (left: *r* = 0.30; right: *r* = 0.30), lateral temporal cortex (left: *r* = 0.50; right: *r* = 0.44), and the temporal poles (left: *r* = 0.21; right: *r* = 0.26). Among subcortical regions, model performance was significantly above chance in the basal ganglia (left: *r* = 0.15, right: *r* = 0.18).

**Figure 3.**
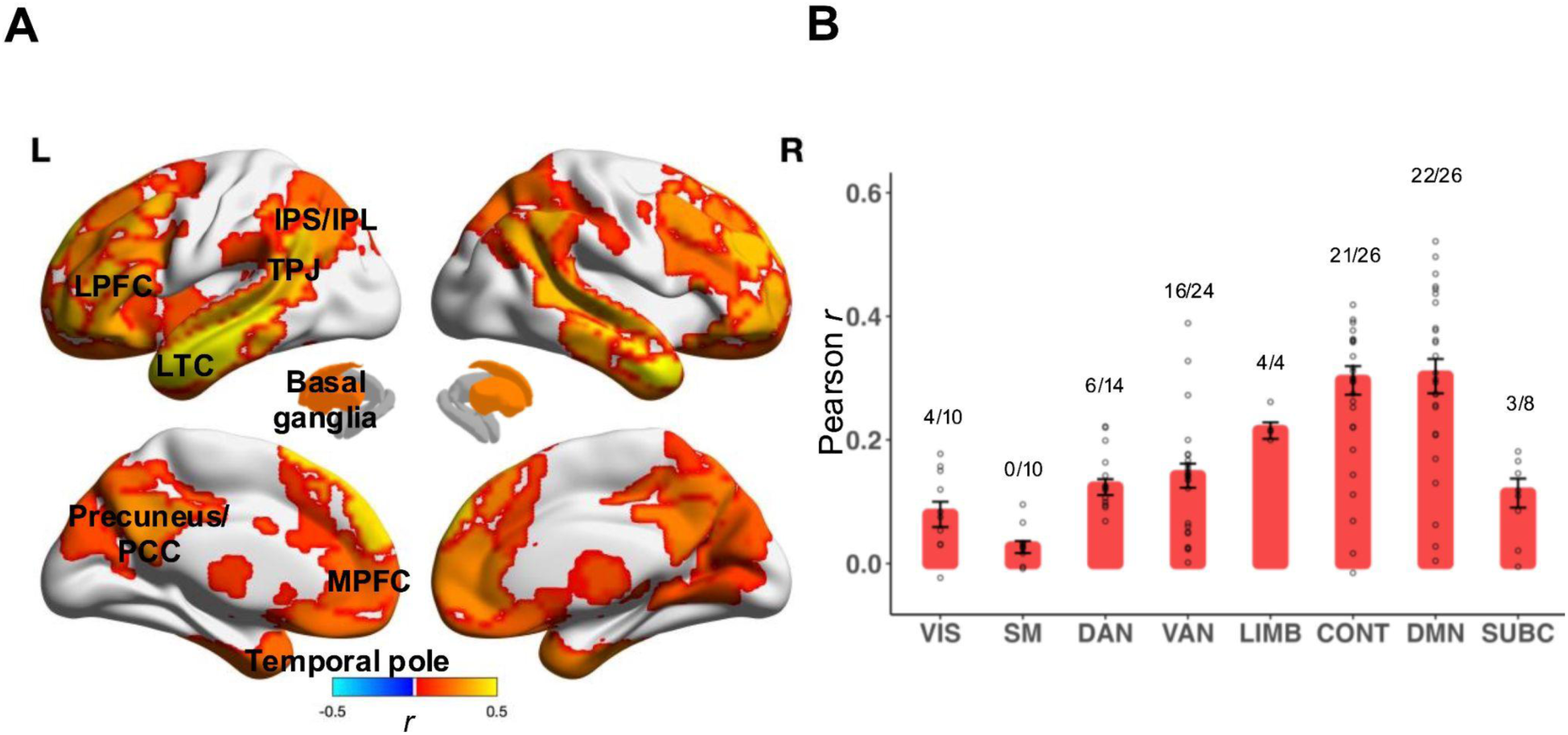
Model performance of fNIRS-fMRI predictive model. **A.** Brain maps showing median correlation between observed BOLD fMRI time courses and predicted BOLD fMRI time courses, thresholded at FDR *q* < 0.05. LPFC: lateral PFC, MPFC: medial PFC, PCC: posterior cingulate cortex, IPS: intraparietal sulcus, IPL: intraparietal lobules; TPJ: temporal parietal junction. **B.** Predictive accuracy by functional network. Height of bar graphs indicates median *r* with circles indicating individual ROIs. Fractions denote the proportion of ROIs significant within a network. See methods for labels of individual networks.

To examine how model performance varies by functional network, we grouped the 122 ROIs into seven cortical functional networks and one subcortical network that includes all the subcortical ROIs. We then calculated the proportion of significant ROIs and median model performance within each network (**Figure 3B**). Model performance was highest in the default mode network (DMN; median *r* = 0.303, percentage significant ROIs = 84.6%) and control network (CONT; median *r* = 0.296, percentage significant ROIs = 80.8%). In contrast, model performance was lowest in the somatosensory network (SM; median *r* = 0.027, percentage significant ROIs = 0%).

### Predicted intersubject functional connectivity patterns from prefrontal fNIRS

To further explore the utility of prefrontal fNIRS data in capturing whole-brain dynamics, we examined whether the predicted fMRI time courses could recapitulate the intersubject functional connectivity (ISFC) patterns observed in the actual fMRI data. ISFC computes functional connectivity across participants and reflects the pattern of whole-brain functional correlations elicited by a shared stimulus (Simony *et al*., 2016). ISFC computed from the observed BOLD was significantly correlated with ISFC computed from the predicted BOLD (*r* = 0.42, *p* < .001, significance assessed using a non-parametric Mantel test; **Figure 4**), indicating that the predicted BOLD successfully reproduced patterns of stimulus-driven functional correlations observed in the actual fMRI data.

**Figure 4.**
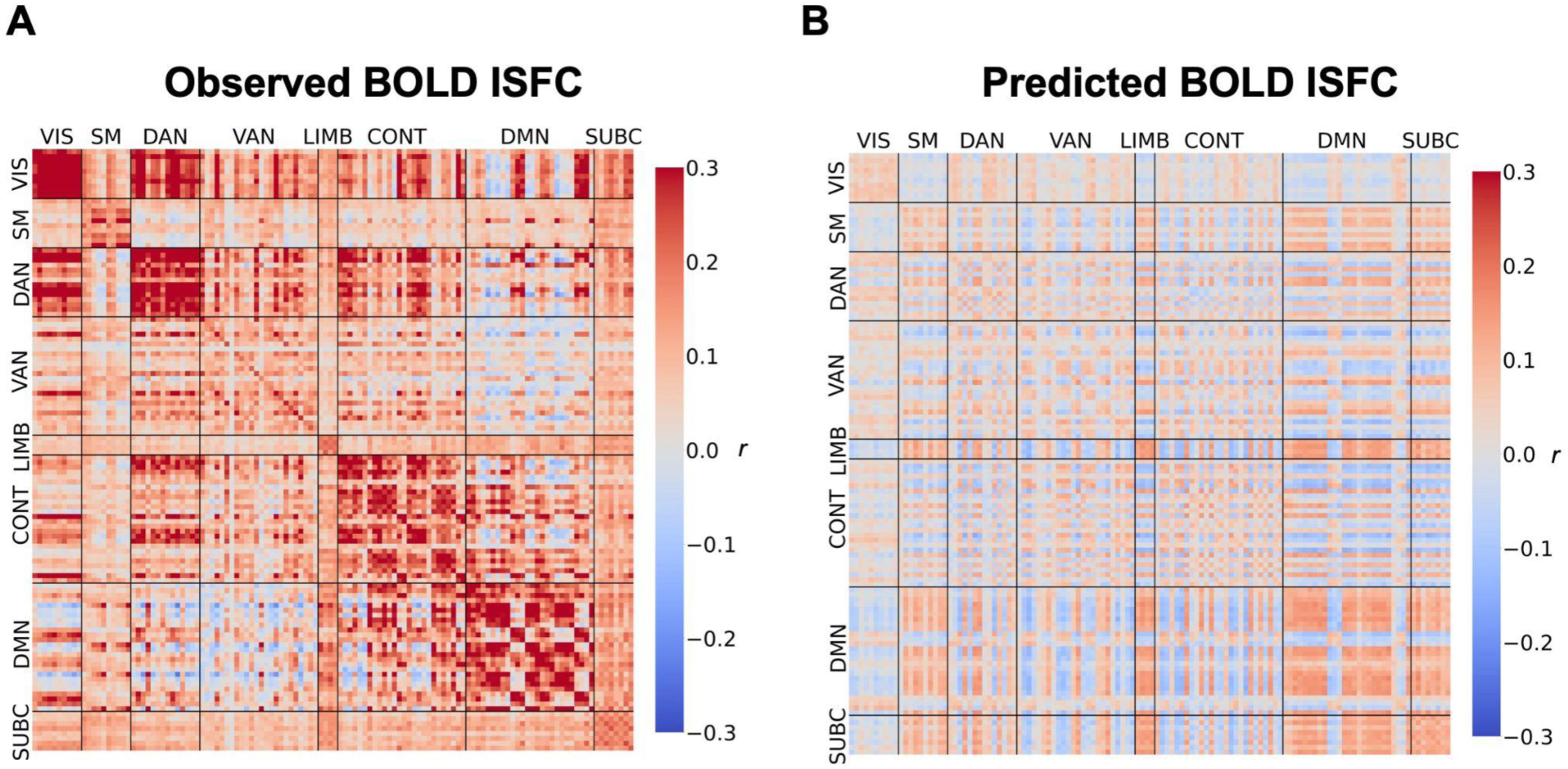
Intersubject functional connectivity (ISFC) matrices computed from A. observed and B. predicted BOLD. Each cell denotes the median ISFC of a pair of ROIs. The correspondence between the two ISFC matrices are computed using a Mantel test. For our analysis, we averaged the top and bottom triangles of the ISFC matrices and excluded the diagonal such that each pair of ROIs is only considered once.

### Assessing semantic information encoded in predicted BOLD

Do the predicted BOLD responses retain semantic information about the movie content? To investigate this, we converted detailed annotations of the movie content into semantic embeddings using the Universal Sentence Encoder (Cer *et al*., 2018; see Methods). We first fit an encoding model to predict the observed Run 1 BOLD time courses from the semantic embedding of each TR. When evaluated on observed BOLD data in Run 2, the encoding model had above-chance accuracy in 51 out of the 66 ROIs where the fNIRS-fMRI model exhibited significant predictive accuracy (FDR *q* < 0.05; **Figure 5A**), indicating that the observed BOLD time courses encode information about the semantic content of the movie.

**Figure 5.**
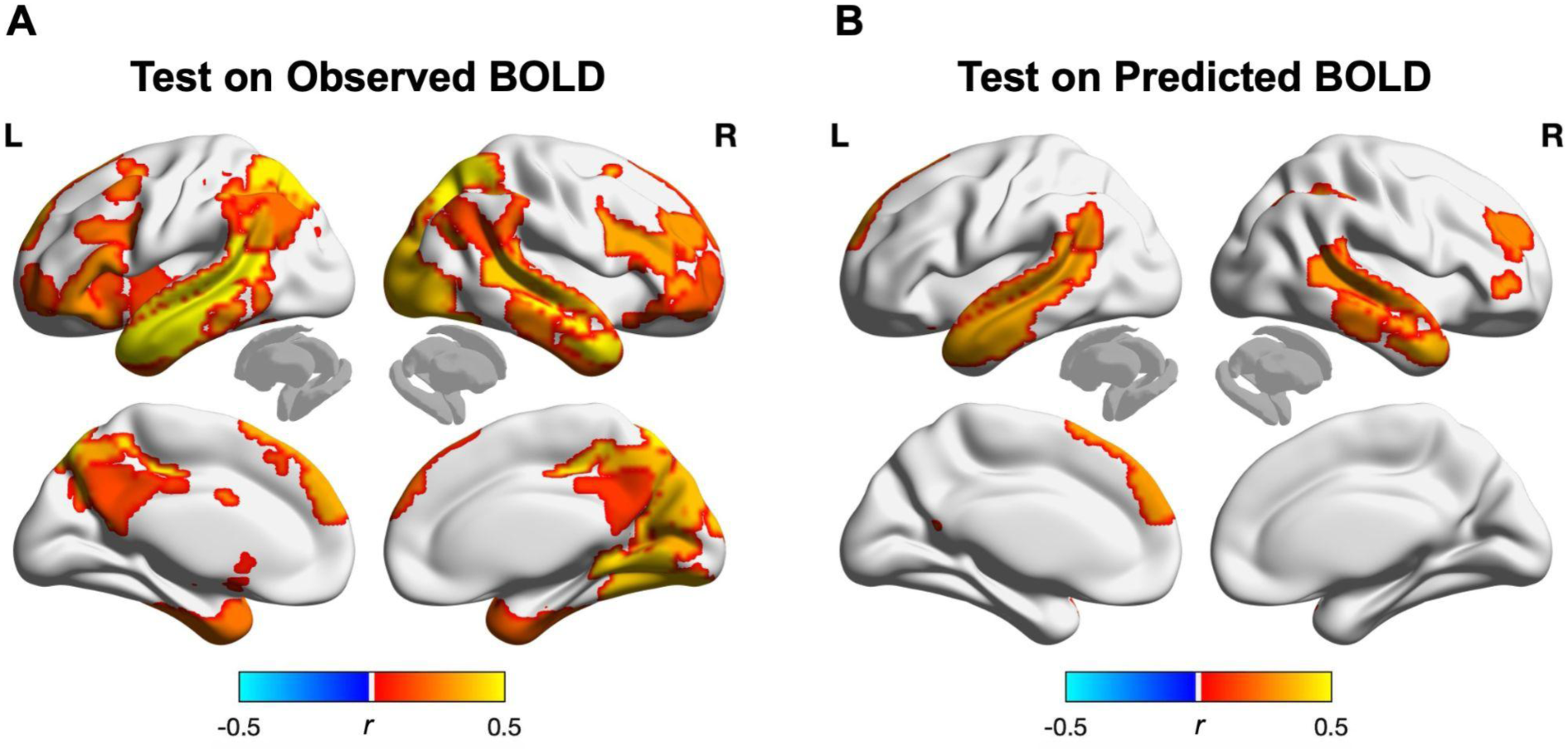
Prediction accuracy of semantic encoding model. An encoding model was trained to predict Run 1 BOLD responses from semantic embeddings of the movie content. **A.** Prediction accuracy of the model on Run 2 observed BOLD responses. **B.** Prediction accuracy of the model on Run 2 predicted BOLD responses. Statistical significance was assessed by retraining the encoding model on phase-randomized training data. Brain maps are thresholded at FDR *q* < 0.05.

Next, to assess whether the predicted BOLD time courses retained this semantic information, we tested the same encoding model on the Run 2 predicted BOLD time courses. Model accuracy was above chance in 12 out of the 66 ROIs (FDR *q* < 0.05; **Figure 5B**), including the dorsomedial prefrontal cortex and the network of brain regions that constitute the language network (i.e., dorsolateral prefrontal cortex, lateral temporal cortex) (Fedorenko *et al*., 2024). These results suggest that the predicted BOLD time courses retained semantic information of the movie content encoded in the original BOLD signal during movie viewing.

## Discussion

In this study, we developed a predictive model that maps prefrontal fNIRS activity to whole-brain fMRI activity while participants viewed naturalistic audio-visual movies. Using this model, we predicted the fMRI activity of participants watching one part of a TV episode based on the fNIRS activity of a different group of participants watching another part of the episode. Our model significantly predicted fMRI time courses above chance in a substantial number of brain regions, including areas not typically accessible by fNIRS, such as the precuneus, temporal poles, and basal ganglia. The predicted fMRI time courses recapitulated intersubject functional connectivity patterns observed in the actual fMRI data, demonstrating the model’s ability to capture whole-brain functional correlations. Furthermore, an encoding model trained to predict observed fMRI data from semantic embeddings showed above-chance accuracy when applied to the predicted fMRI time courses generated by our model, suggesting that the model-generated fMRI time courses retained semantic information encoded in fMRI activity. Overall, our findings demonstrate the feasibility of using fNIRS to infer broader brain activity patterns, opening up new possibilities for investigating brain function during complex, real-world contexts.

fNIRS relies on measurements of changes in light absorption near the surface of the scalp. To relate these measurements to brain function, we need to map these changes onto neural activity. Existing methods have primarily focused on anatomical correspondence (Tsuzuki and Dan, 2014) where fNIRS measurements are mapped onto cortical areas between sources and detectors. While useful, this anatomically constrained approach limits fNIRS-based inferences to regions directly beneath the optodes and relies heavily on optode placement being consistent across participants. In contrast, our study takes a complementary approach by leveraging patterns of co-activation during tasks across fNIRS and fMRI (Liu *et al*., 2015). Using this approach, we show that fNIRS signals can not only be mapped to their anatomically corresponding regions but also used to predict neural dynamics in functionally interconnected regions.

Our study measured fNIRS activity only from the prefrontal cortex. Depending on the number of optodes available, researchers do not always have the number of optodes to achieve coverage across the entire scalp. This is especially so in hyperscanning studies (Hamilton, 2021) where a researcher might want to split up the optodes to measure neural activity across multiple participants. The PFC is an ideal target due to its functional heterogeneity and diverse connections to multiple large-scale networks that can be leveraged to predict widespread brain activity (Menon and D’Esposito, 2022). Additionally, the PFC often provides cleaner fNIRS signals because it is less obstructed by hair compared to other scalp regions. Our study provides proof of the concept that even with limited coverage, one can predict neural activity in approximately half of the brain, including in areas that were anatomically inaccessible by fNIRS.

Model accuracy was highest in the default mode network and control network, both of which are critically involved in many higher-order cognitive functions (Raichle, 2015; Gratton *et al*., 2018; Menon and D’Esposito, 2022). This suggests that the current model may be particularly well suited for investigating cognitive processes that rely on these networks, including social cognition, cognitive control, working memory, introspection and memory retrieval. Conversely, model prediction accuracy was lowest in the somatomotor network. This discrepancy is likely due to the nature of the task we employed rather than a limitation of the approach. Our model was trained on data collected while participants passively watched movies, which have been previously found to engage both the default mode network and the control network (Simony *et al*., 2016; Song *et al*., 2021; Yeshurun *et al*., 2021), thus allowing our model to learn mappings for these regions. Watching a movie, however, does not involve tactile stimulation or movement, and is thus unlikely to engage somatomotor areas. As a result, the model had limited training data to learn robust mappings for these regions. Future work can potentially expand the coverage of our model by incorporating paradigms that engage the somatomotor network (e.g., playing a musical instrument).

It is important to note that our approach approximates neural activity based on functional correlations, and highly connected areas may serve related but distinct functions. For example, the dorsomedial PFC and temporoparietal junction are both part of the default mode network and are often co-activated during social interaction, but may support distinct processes (Saxe, 2006; Van Overwalle, 2009; Konovalov *et al*., 2021). At present, it is unclear whether our model captures the level of granularity needed to differentiate between the distinct roles of such interconnected regions. Instead, we see the utility of our model as a tool for generating hypotheses about broader patterns of brain activity that can guide future investigations with fMRI. By identifying large-scale functional relationships and co-activation patterns, our approach can highlight regions or network interactions that warrant deeper exploration using imaging techniques with higher spatial resolution. This iterative process - using fNIRS-based predictions to inform targeted fMRI studies can help uncover more nuanced brain functions and advance our understanding of the distinct roles played by interconnected regions.

Our approach shares similarities with functional alignment techniques, such as hyperalignment (Haxby *et al*., 2011; Haxby *et al*., 2020) and the Shared Response Model (SRM; Chen *et al*., 2015). These methods aim to align neural responses across individuals to a common representational space via a shared stimulus, facilitating the comparison of brain activity across different participants. Our fNIRS-fMRI predictive model is similar to these approaches in that we leverage the shared neural dynamics elicited by naturalistic stimuli to align neural responses. However, our approach is unique in that it aligns responses collected using different neuroimaging modalities, each with different spatial resolutions and coverage. Using this approach, we were able to predict the average activity time course of participants watching an unseen stimulus based on the fNIRS data from a different participant, demonstrating cross-participant and cross-stimulus generalizability. We note that our study used two parts of the same TV episode to train and test the model, and the inherent similarities in the content between the two parts may help boost model performance. Future work applying the model on entirely different stimuli will provide a more stringent test of the model’s generalizability across context.

Similar to other functional alignment approaches that rely on shared stimulus-driven responses, our model primarily captures responses that are consistent across individuals. However, some brain regions tend to exhibit more idiosyncratic responses even when processing the same stimulus. For example, prior fMRI work had found that ventromedial prefrontal cortex (VMPFC) responses tend to be idiosyncratic when participants watched a TV episode (Chang *et al*., 2021). Similarly, we had previously shown that fNIRS responses in the VMPFC while listening to social situations were modulated by individual differences in social beliefs (Lyu *et al*., 2024). In line with these findings, the accuracy of our fNIRS-fMRI model was lower in the ventromedial prefrontal regions relative to dorsolateral or dorsomedial prefrontal regions, suggesting that the model was less able to capture the more idiosyncratic responses in the VMPFC. Measuring the same person watching the same video using both fNIRS and fMRI may allow us to build individual-specific models. These personalized models could then be used to predict fMRI activity from fNIRS data of the participant performing tasks that are not amenable to fMRI. If this approach proves effective, incorporating a movie stimulus into fMRI scans could be a worthwhile strategy to allow for the alignment of an individual’s brain to other modalities more broadly.

To examine the information content encoded in the predicted fMRI signal, we built on a growing body of research utilizing text embedding models to study how semantic information is encoded in the brain (e.g., Huth *et al*., 2016; Pereira *et al*., 2018; Goldstein *et al*., 2022; Caucheteux *et al*., 2023). Embedding models convert text (i.e., words or sentences) into high-dimensional vector representations that reflect their semantic meaning. Findings from these studies have shown that these semantic embeddings can be mapped onto brain activity, suggesting that the brain’s response to semantic content can be quantitatively modeled using these models. In our study, we first replicated these findings by demonstrating that an encoding model trained to predict BOLD time courses from semantic embeddings of the movie content exhibited above-chance accuracy, indicating that the BOLD time courses captured information about the semantic content of the movie.

Importantly, the same encoding model showed above-chance accuracy when applied to the BOLD time courses generated by our fNIRS-fMRI model, with prediction accuracy highest in the dorsolateral PFC and lateral temporal cortex. Both regions are part of a language network that supports language comprehension and production (Fedorenko *et al*., 2024). Significant prediction was also observed in the dorsomedial PFC, a region involved in the interpretation of narratives (Yeshurun *et al*., 2017; Nguyen *et al*., 2019; Leong *et al*., 2020). The high accuracy of the encoding model in these areas is consistent with the notion that these regions are involved in the processing and integration of semantic information. Moreover, these results suggest that the predicted BOLD retains content-specific information about participants’ cognitive experiences, highlighting the potential of using our approach to study how the brain encodes complex, naturalistic experiences in real-world contexts where fMRI is not feasible.

The overarching objective of our project is to develop a tool that enhances the versatility of fNIRS. To that end, we have made an fNIRS-fMRI model trained on all of the Run 1 data publicly available at https://github.com/ycleong/fNIRS-fMRI_models, as well as example scripts on how to load and apply the model. We note that unlike the analyses presented in this paper, where the model was trained using a leave-one-participant-out approach to assess cross-participant generalizability, this version of the model was trained on Run 1 data of all participants to provide a single model for use. We are committed to the continued development of this tool, including training and validating our models on additional stimuli and tasks, as well as developing models for different fNIRS montages. We also invite the fNIRS community to utilize and contribute to this tool, with the goal of enhancing the utility of fNIRS in diverse research contexts and facilitating its application in studying complex, real-world behaviors.

## Data Availability Statement

Preprocessed MRI data from the *Sherlock* dataset can be accessed at Princeton’s DataSpace, https://dataspace.princeton.edu/jspui/handle/88435/dsp01nz8062179. Preprocessed fNIRS data can be accessed at https://github.com/ycleong/fNIRS-fMRI_models.

## Acknowledgements

This research was supported by computing resources provided by the University of Chicago Social Sciences Division, the University of Chicago Research Computing Center, and the University of Chicago Data Science Institute. We would also like to thank the University of Chicago Neuroscience Institute Shared Equipment Award for providing the fNIRS device.

## Author Contributions

SG and YCL contributed to conceptualization and methodology. SG served as a lead for investigation, data curation, formal analysis, software, validation, visualization, and writing — original draft. RN served in a supporting role for validation, visualization, formal analysis, and software. SB contributed to software. SG, RN, and YCL contributed equally to writing — review & editing; SB served in a supporting role for writing — review & editing. YCL contributed to supervision, resources, and project administration.

