## Supplementary Information for "Predicting whole-brain neural dynamics from prefrontal cortex fNIRS signal during movie-watching"

**Table S1. Table showing the correlation between the 20 fNIRS channels and matching fMRI ROIs.** **: $q < 0.01$; *$q < 0.05$.

| Channel | ROI | r-value | p-value | q-value |
|---|---|---|---|---|
| **1** | left dorsolateral PFC | 0.116 | 0.011 | 0.015* |
| **2** | left lateral PFC | 0.075 | 0.039 | 0.0433* |
| **3** | left ventrolateral PFC | 0.242 | 0.001 | 0.002** |
| **4** | left orbitofrontal PFC | 0.283 | 0.001 | 0.002** |
| **5** | left dorsolateral PFC | 0.207 | 0.001 | 0.002** |
| **6** | left frontopolar cortex | 0.179 | 0.001 | 0.002** |
| **7** | left medial PFC | 0.513 | 0.001 | 0.002** |
| **8** | left dorsomedial PFC | 0.111 | 0.026 | 0.031* |
| **9** | dorsomedial PFC | 0.197 | 0.001 | 0.002** |
| **10** | right dorsomedial PFC | 0.077 | 0.082 | 0.086 |
| **11** | left ventromedial PFC | 0.058 | 0.154 | 0.154 |
| **12** | medial PFC | 0.216 | 0.002 | 0.003** |
| **13** | right ventromedial PFC | 0.225 | 0.001 | 0.002** |
| **14** | right medial PFC | 0.274 | 0.001 | 0.002** |
| **15** | right dorsolateral PFC | 0.225 | 0.002 | 0.003** |
| **16** | right frontopolar cortex | 0.201 | 0.002 | 0.003** |
| **17** | right lateral PFC | 0.104 | 0.024 | 0.030* |
| **18** | right dorsolateral PFC | 0.157 | 0.004 | 0.006** |
| **19** | right orbitofrontal PFC | 0.255 | 0.001 | 0.002** |
| **20** | right ventrolateral PFC | 0.227 | 0.001 | 0.002** |

**Table S2. Table showing correlation between fNIRS channel and average A1, V1, and global fMRI signal.** Correction for FDR was performed separately for A1, V1 and global signal.

| Channel | A1 | V1 | Global |
|---|---|---|---|
| **1** | r = -0.053, p = 0.846, q = 0.846 | r = 0.007, p = 0.436, q = 0.712 | r = -0.073, p = 0.891, q = 1.000 |
| **2** | r = -0.027, p = 0.716, q = 0.835 | r = 0.072, p = 0.076, q = 0.304 | r = 0.049, p = 0.167, q = 0.477 |
| **3** | r = -0.036, p = 0.751, q = 0.835 | r = 0.046, p = 0.199, q = 0.497 | r = -0.112, p = 0.975, q = 1.000 |
| **4** | r = 0.044, p = 0.223, q = 0.555 | r = 0.078, p = 0.048, q = 0.304 | r = 0.029, p = 0.268, q = 0.536 |
| **5** | r = -0.037, p = 0.794, q = 0.836 | r = -0.070, p = 0.919, q = 0.996 | r = 0.061, p = 0.125, q = 0.477 |
| **6** | r = 0.079, p = 0.066, q = 0.555 | r = 0.075, p = 0.068, q = 0.304 | r = 0.039, p = 0.217, q = 0.482 |
| **7** | r = 0.062, p = 0.129, q = 0.555 | r = -0.110, p = 0.981, q = 0.996 | r = 0.026, p = 0.321, q = 0.583 |
| **8** | r = 0.016, p = 0.362, q = 0.723 | r = -0.020, p = 0.672, q = 0.961 | r = 0.072, p = 0.045, q = 0.300 |
| **9** | r = 0.044, p = 0.187, q = 0.555 | r = -0.092, p = 0.983, q = 0.996 | r = 0.061, p = 0.094, q = 0.470 |
| **10** | r = 0.031, p = 0.213, q = 0.555 | r = 0.025, p = 0.262, q = 0.505 | r = 0.096, p = 0.016, q = 0.160 |
| **11** | r = 0.075, p = 0.103, q = 0.555 | r = -0.088, p = 0.965, q = 0.996 | r = -0.164, p = 1.000, q = 1.000 |
| **12** | r = 0.008, p = 0.457, q = 0.755 | r = -0.139, p = 0.996, q = 0.996 | r = -0.066, p = 0.891, q = 1.000 |
| **13** | r = 0.035, p = 0.250, q = 0.555 | r = 0.066, p = 0.106, q = 0.328 | r = -0.010, p = 0.583, q = 0.898 |
| **14** | r = -0.030, p = 0.739, q = 0.835 | r = -0.064, p = 0.895, q = 0.996 | r = 0.050, p = 0.158, q = 0.477 |
| **15** | r = -0.024, p = 0.704, q = 0.835 | r = 0.005, p = 0.463, q = 0.712 | r = 0.117, p = 0.003, q = 0.060 |
| **16** | r = 0.045, p = 0.189, q = 0.555 | r = 0.032, p = 0.264, q = 0.505 | r = 0.010, p = 0.423, q = 0.704 |
| **17** | r = 0.001, p = 0.491, q = 0.755 | r = 0.024, p = 0.278, q = 0.505 | r = 0.034, p = 0.210, q = 0.482 |
| **18** | r = 0.049, p = 0.183, q = 0.555 | r = 0.062, p = 0.115, q = 0.328 | r = -0.061, p = 0.853, q = 1.000 |
| **19** | r = -0.021, p = 0.697, q = 0.835 | r = 0.087, p = 0.055, q = 0.304 | r = -0.024, p = 0.683, q = 0.976 |
| **20** | r = 0.004, p = 0.482, q = 0.755 | r = 0.077, p = 0.070, q = 0.304 | r = -0.109, p = 0.963, q = 1.000 |
